# Genetically encoded split luciferase biosensors to measure endosome disruption in real time in live cells

**DOI:** 10.1101/2020.01.16.906180

**Authors:** Kameron V. Kilchrist, J. William Tierney, Craig L. Duvall

## Abstract

Endosomal escape is a critical step in intracellular delivery of biomacromolecular drugs, but quantitative, high throughput study of endosomal vesicle disruption remains elusive. We designed two genetically encoded split luciferase “turn on” reporters that can be assayed rapidly in well plates on live cells using a luminometer. Both systems use non-luminescent N-terminal and C-terminal luciferase fragments which can reconstitute a functional luminescent enzyme when they are held in proximity by their fusion partners. The first system uses Gal8 and CALCOCO2 fused to these fragments, which interact following endosome disruption and facilitate complementation of the split luciferase fragments to produce significant luminescence when luciferin is added. The second system uses the N-terminal carbohydrate recognition domain of Gal8 (G8-NCRD) fused to both luciferase fragments. Following endosome disruption, G8-NCRD binds to exposed glycans inside endosomes, concentrating both fragments there to reconstitute active luciferase. Additionally, and in contrast to recently reported Gal8 intracellular tracking with fluorescent microscopy, these split luciferase-based assays enable simultaneous identification and downselection of cytotoxic test conditions because the luciferase reaction requires intracellular ATP. Further, we demonstrate that the lead luminescent cell line is more sensitive to detection of endosomal disruption at lower doses of an endosome disrupting drug carrier than the previously reported Gal8-YFP fluorescent system. These systems represent a first-in-class luminescent assay to detect endosome disruption in high throughput while excluding toxic formulations. Endosome disruption screening with these “turn on” systems has potential as a tool in the discovery and development of intracellular biologic drug delivery formulations.

**Graphical Abstract:** 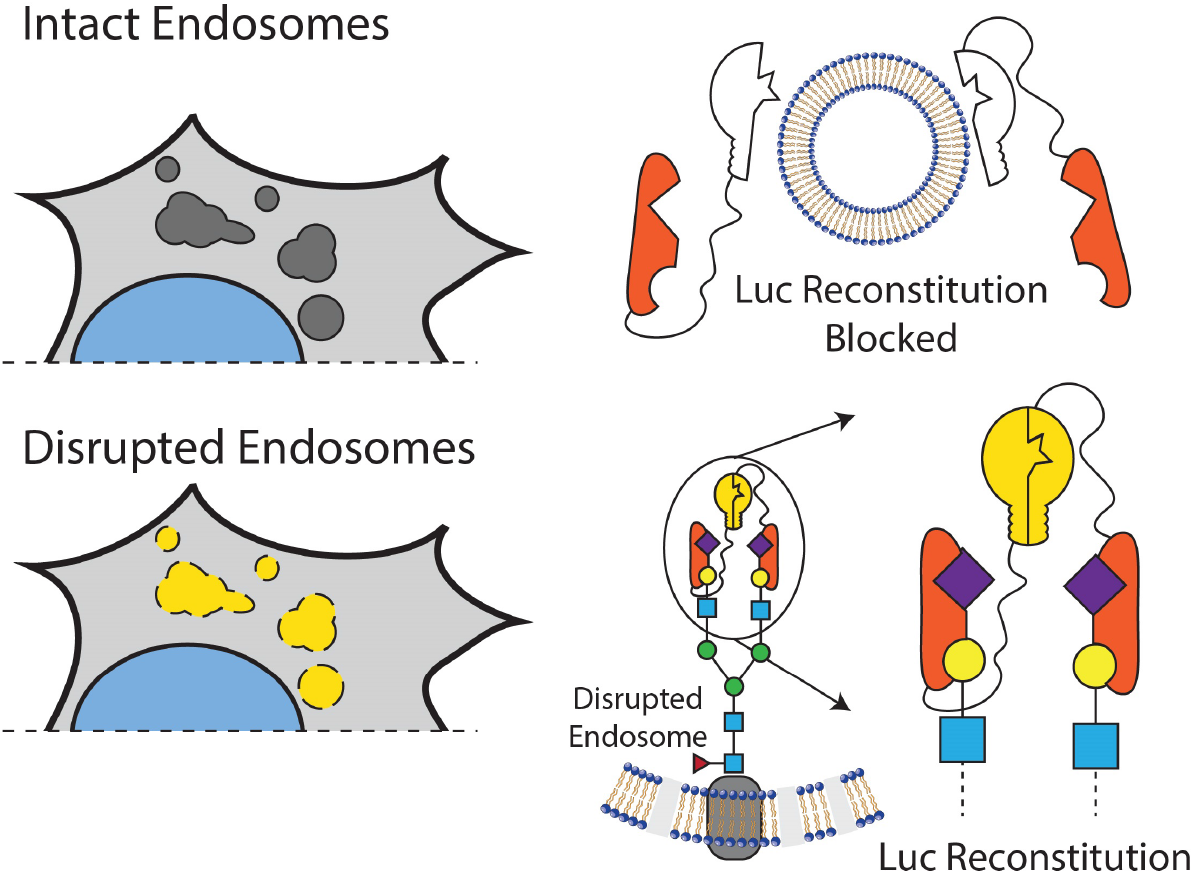

## Introduction

The intracellular endosomal membrane remains a critical barrier to large molecule drugs with intracellular mechanisms of action. While there are now several exciting FDA-approved large molecule drugs which act inside the cell, including an siRNA lipid complex nanomedicine which uses active endosome-disruption technology^1–3^, the broad-scale clinical utilization of intracellular-acting biologic medicines has been hindered by challenging pharmacokinetics in the circulatory system, specific tissue-level barriers to delivery^4,5^, and barriers at the cellular and subcellular level. At the cellular level, large, hydrophilic molecules such as nucleic acids and peptides are internalized by endocytosis, which compartmentalizes them away from cytosolic drug targets and normally leads to lysosomal trafficking and degradation. A common strategy to promote cell entry and endosomal escape is formulation into nanoscale drug delivery systems, which are often engineered to have pH-responsiveness that is activated by endosome acidification to promote endosomal membrane disruption and translocation.

While endosome disruption is an important feature for nanoscale drug delivery systems, this process is difficult to measure rapidly and quantitatively. Several methods, including subcellular fractionation^6^ and transmission electron microscopy,^7^ can be used to assess cytosolic distribution and endosomal disruption in a low throughput format. We and others have recently investigated tracking of intracellular galectins as a method to assess endosomal integrity^8^ in live cells^9–12^; this method has been applied by us and others to assess endosome disrupting nanomedicine formulations.^9–20^ Galectins are a family of carbohydrate binding proteins which serve to mediate cell-cell and cell-matrix interactions, modulate immune cell functions, and restrict intracellular pathogens.^21^ Glycosylation on the inner leaflet of endosomes becomes accessible to cytosolic Galectin-3, -8, and -9, which bind and concentrate there following membrane disruption by pathogens^8^ or drug carriers.^9–11^ By quantitative tracking of fluorescence redistribution from diffuse (in the cytosol) to punctate (inside disrupted endosomes), the extent of endosomal disruption by pathogens and drug carriers can be assessed.

We have been particularly interested in the development of Gal8 based assays to monitor endosomal disruption in the context of subcellular trafficking of drug carriers.^9–11^ Galectin 8 (Gal8) is specifically involved in the detection of endosomal disruption and is critical to the restriction of intracellular infection by pathogens.^8^ Gal8 is a tandem-repeat lectin, containing two carbohydrate recognition domains (CRD) at the N and C termini with dissimilar substrate specificities. The N-terminal CRD (G8-NCRD) binds to host rather than pathogen glycans^8^, specifically to sialylated glycans containing α2-3-sialylated or 3′ sulfated β-galactosides.^22^ Meanwhile, the C-terminal CRD mediates trafficking of damaged endosomes to an autophagy pathway by interacting with the C terminus of CALCOCO2, which recruits the adapter protein LC3 to induce engulfment of the damaged vesicle (Figure 1A).^8,9,23^

**Figure 1.**
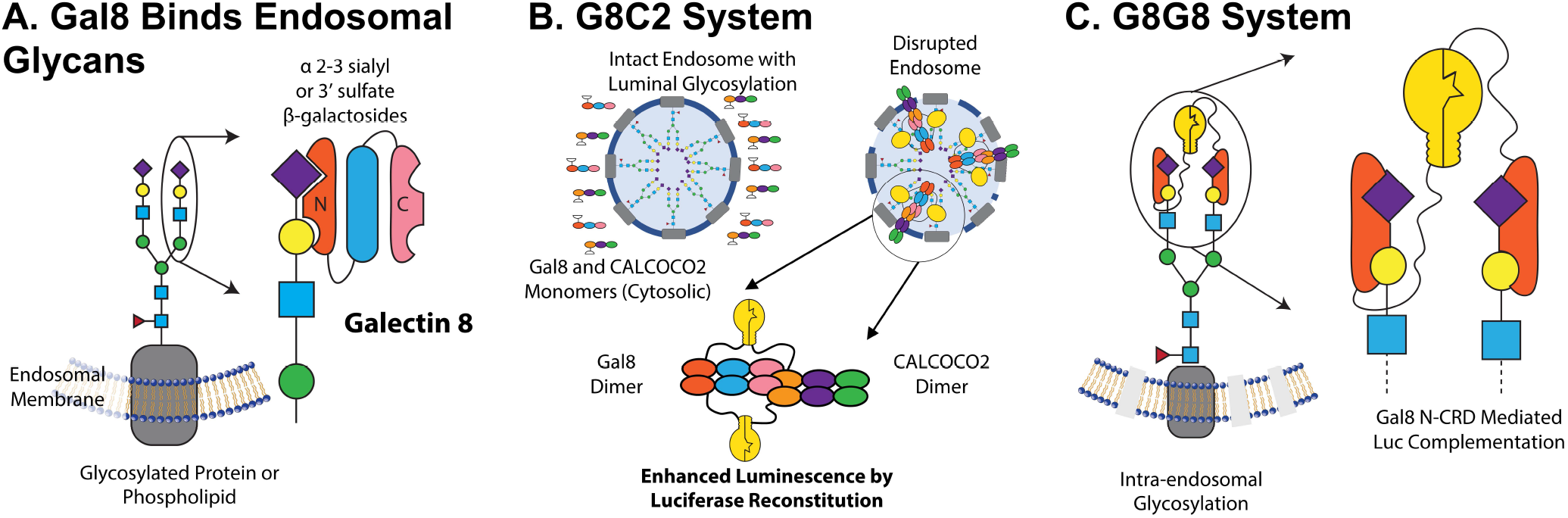
Overview of Gal8 activity and engineered endosome disruption reporters. (A) Gal8 binds to intra-endosomal α 2-3 sialyl or 3′ sulfated β-glycans through its N terminal carbohydrate recognition domain (CRD) upon disruption of the endosomal membrane. (B) Following endosomal disruption, Gal8 binds to intraendosomal glycans and recruits CALCOCO2 through a dimer-dimer complex to initiate macroautophagy of the damaged vesicle. The G8C2 system was designed to detect this protein-protein interaction, with full length Gal8 constructed as an N-terminal fusion to the N-terminal luciferase fragment and full length CALCOCO2 as a C-terminal fusion to the C-terminal luciferase fragment. (C) The G8G8 system uses the specific binding of Gal8 N-terminal CRD to concentrate and facilitate reconstitution of NLuc and CLuc fragments on the interior leaflet of the disrupted endosomal membrane. The N-terminal and C-terminal luciferase fragments were fused to the N-terminus and C-terminus, respectively, of the Gal8 N-CRD fragment.

We previously demonstrated a method of measuring endosomal disruption by quantifying the total recruited Gal8 on a per cell basis. We showed that this method correlated to cytosolic siRNA bioavailability and that it was more predictive of cytosolic drug bioactivity than methods traditionally used to measured endosome disruption.^11^ In *in vivo* screening, Gal8 tracking in MDA-MB-231 orthotopic tumors extracted from mice after treatment with an endosomolytic nanocarrier showed statistically significant endosomal disruption compared to vehicle control injections. However, in cells, this method required high content microscopy capable of screening thousands of cells for robust analysis, while in tissue, subcellullar Gal8 redistribution was difficult to image *ex vivo*. Further, because subcellular imaging through the skin is difficult, tumors were excised from the mice for imaging, precluding the ability to perform kinetic analyses. To overcome these challenges, we sought a turn on style assay with faster data acquisition, the potential for simple and longitudinal intravital measurements, and the capability to be implemented without requiring advanced microscopy or computational image analysis.

In the current work, we developed a turn on style endosome disruption assay based on a split luciferase system, wherein the 550 amino acid firefly luciferase is split into two overlapping fragments (A.A. 1-398 and 394-550). This split luciferase system was chosen because it is a low affinity pair suitable for detecting transient protein-protein interactions by producing a significant increase in luminescent signal when the two fragments of the enzyme are brought into close proximity, facilitating their complementation.^24^ The use of a luciferase based system, as opposed to a fluorescent reporter, was attractive because firefly luciferase is widely used *in vitro* and *in vivo* and utilizes the inexpensive and nontoxic substrate D-luciferin. Firefly luciferase signal (unlike *renilla* or related coelenterazine-dependent luciferases) also requires ATP, which is released from cells upon loss of viability, reducing the chances of identifying cytotoxic formulations as false positive assay hits.

We designed two split luciferase reconstitution systems to test as reporters for high throughput measurement of endosomal disruption. The first design, called G8C2, was constructed to measure the protein-protein interactions of full length copies of Gal8 and CALCOCO2 (also known as NDP52), which have been shown to interact downstream of Gal8 clustering to initiate LC3-mediated macroautophagy of damaged endosomes.^8^ Full length copies of Gal8 and CALCOCO2 were connected N-terminal to human codon optimized Luc2 amino acids 1-398 (NLuc398) and C-terminal to luciferase amino acids 394-550 (CLuc394), respectively, by a 3x(GGGGS) linker. An internal ribosomal entry site was used to drive expression of both partners from the same mRNA transcript (Figure S1 A). Our design was informed by the recently-elucidated structure of the Gal8/CALCOCO2 interaction, such that NLuc398 and CLuc394 would be in close proximity upon the dimer-dimer interaction of Gal8/Gal8 and CALCOCO2/CALCOCO2 complexes (Figure 1B).^23^ We hypothesized that such a system would demonstrate enhanced luminescence upon endosomal disruption due to the purported interaction between Gal8 and CALCOCO2.

The second reporter design, called G8G8, was based on the hypothesis that a single carbohydrate binding domain from Gal8, G8-NCRD, was sufficient to drive complementation of NLuc and CLuc fragments. NLuc398 was connected as an N terminal fusion to G8-NCRD, while CLuc394 was connected as a C terminal fusion to G8-NCRD, both using a 3x(GGGGS) linker (Figure S1 B). Upon intra-endosomal concentration on glycans, NLuc and CLuc fragments could reconstitute and produce luciferase signal on the endosomal glycocalyx. A schematic of the anticipated interaction is shown as Figure 1C.

To assess the suitability of these systems to monitor endosomal escape, we designed a single lentiviral transfer vector for each system, generated lentiviral particles, and created five stable cell populations using a dilution series of virus. We then assessed luciferase signal turn on in response to treatment with PPAA, a cytocompatible and robust inducer of endosomal disruption. We then used the lead cell population to validate endosome disruption from three commercial reagents: the commercial cationic lipids 1,2-dioleoyl-3-trimethylammonium-propane (DOTAP) and Lipofectamine^®^ 2000, as well as the *in vivo* transfection reagent JetPEI^®^. We also cross-validated with four members of our cationic diblock copolymer molecular weight library which have been previously assessed with the Gal8-YFP method and produce varying levels of endosome disruptive potential that correlate with polymer molecular weight.^11,25^ These polymers, known as “50B,” are diblock structures of PEG and a random copolymer of 50 mol% of the basic monomer dimethylaminoethyl methacrylate (DMAEMA) and 50 mol% of the hydrophobic monomer butyl methacrylate (BMA).

## Results and Discussion

### Response of G8C2 and G8G8 Systems to PPAA

To induce endosomal disruption, we used the anionic pH-responsive polymer poly(propylacrylic acid) (PPAA), a well-studied intracellular delivery reagent,^10,15,26–30^ that we have previously shown induces rapid endosomal disruption detectable using the microscopy-based Gal8-YFP assay within 30 minutes.^10,15,16^ We tested five different populations of HEK293-T cells generated from a dilution series of G8C2 and G8G8 system encoding lentivirus to assess the relative effect of transduction ratio on luminescent response due to PPAA treatment (Figure S2 and S3, left). In this initial screen, two cell populations expressing the G8C2 system and all five cell populations expressing the G8G8 system produced statistically significant dose responses to PPAA treatment. After normalizing the luminescence data to the average of buffer controls, we plotted fold-response curves (Figure S2 and S3, right). The G8C2 system transfected at the highest multiplicity of infection had the highest overall fold-change response (Figure 2A and S2), while all G8G8 cell populations produced significant response to PPAA. Furthermore, all G8G8 cell populations except one show response to a wide range of PPAA doses (Figure S3). It is also observable from these data that PPAA induces cytotoxicity at 1250 and 625 μg/mL dosing (Figure 2A and 2B), which is detected as luminescence signal below baseline (vehicle treated cells). Cytotoxicity in this dose range was also cross-validated with the resazurin cytotoxicity assay (Figure S4). Interestingly, the G8G8 system produced an order of magnitude brighter overall luminescence (Figure 2B) than the G8C2 system. This likely contributed to the lower variance of measured signal, improving statistical confidence of measurements using the G8G8 assay. This higher signal and lower variance may also prove useful in the application of this system as an *in vivo* reporter.

**Figure 2.**
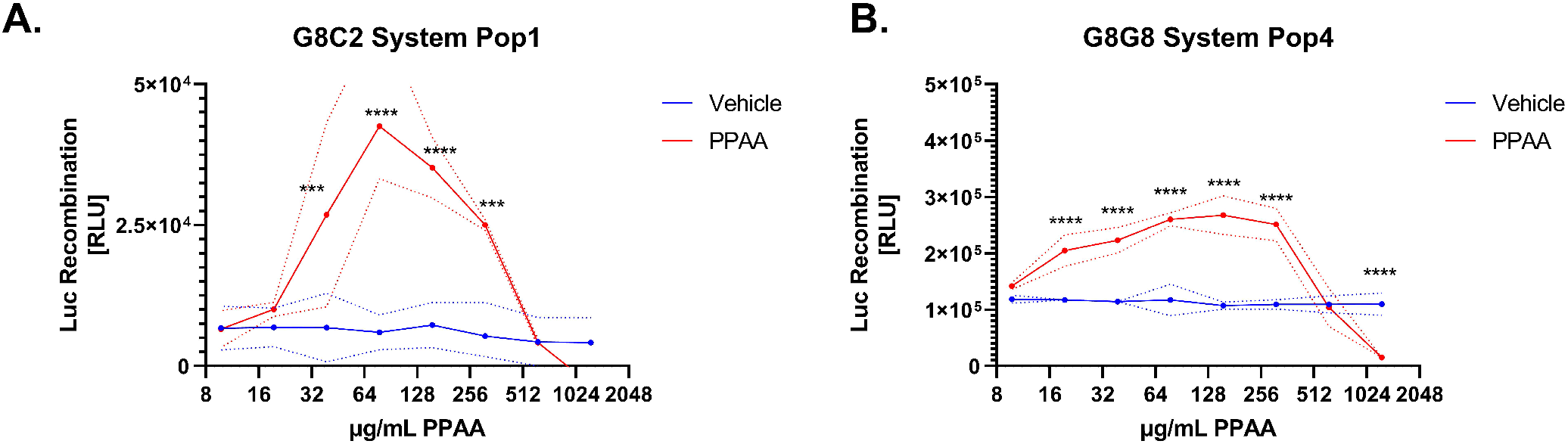
Lead candidate G8C2 and G8G8 cell populations demonstrate robust luciferase reconstitution following endosomal disruption. G8C2 and G8G8 HEK293-T cells were incubated with a 2-fold dilution series from 1250 to 9.8 μg/mL PPAA for 2 hours. Both systems show cell toxicity at the two highest doses tested (1250 and 625 μg/mL), visible as luminescence at or below vehicle-treated cell baseline. Both systems also generate significant luminescent response at doses from 313 to 39 μg/mL for G8C2 population 1 (Pop1) and from 313 to 20 μg/mL for G8G8 population 4 (Pop4). Additional information on statistical tests is provided in supplementary methods, and full statistical test results are reported in Supplementary Tables 1 and 2.

We also calculated the area under the curve (in units of fold-response × μg/mL) to assess both the G8G8 and G8C2 systems (Figure S5). This analysis revealed that AUC correlated to viral concentration during transduction for the G8C2 system, with a statistically significant positive nonzero slope (p < 0.05, R^2^ = 0.90); these data indicate that high transduction ratios are advantageous for the G8C2 system.

Making a series of cell populations with varied multiplicities of infection was initially motivated by the idea that luciferase complementation may occur and create baseline signal (without co-concentration of the reporter proteins into disrupted endosomes) if the reporter proteins were overexpressed to too high of a level. However, higher multiplicity of infection correlated with better readouts with the G8C2 system. In this system, the interaction between NLuc-Gal8 and CALCOCO2-CLuc is in competition with their respective binding partners being endogenously produced within the cell, as it is expected that the split luciferase tagged proteins also form non-luminescent complexes with the endogenous protein binding partners. However, the G8G8 split luciferase system responsivity was not sensitive to transduction ratio. In the G8G8 system, both Gal8-CLuc and Gal8-NLuc, as well as endogenous Gal8, are competing for binding to glycans that cover the internal leaflet of the endosomal membrane. We hypothesize that the competition with endogenous Gal8 in this system may be less impactful because the luminal surface of the endosome provides a larger reaction surface area available to coordinate luciferase reconstitution, versus the fixed conformation of the dimer-dimer interaction of CALCOCO2 and Gal8.

Another potential advantage of the G8G8 system is that it uses a minimal N-terminal protein domain derived from Gal8 in the reporter, which is in opposition to the G8C2 system that expresses full-length Gal8 and CALCOCO2. As a result, the G8G8 design uses a shorter DNA sequence (3,318 base pairs for G8G8 vs. 4,344 base pairs for the G8C2 system), which allows additional elements to be introduced into the same lentiviral vector (which are limited to approximately 6,400 base pairs). For example, here we additionally incorporated an EGFP element to assess transduction efficiency by microscopy into the G8G8 system. Use of the minimal N-terminal Gal8 system may also be less disruptive to normal cellular processes, as the effector functions of Gal8 signaling as an endosomal damage sensor occur through its C-terminal carbohydrate recognition domain^23^, which is not expressed by our G8G8 construct. For these reasons, we focused our subsequent, deeper characterization and benchmarking on the G8G8 system.

### Comparisons to Validated Gal8-YFP Endosomal Escape Assay

A HEK293-T Gal8-YFP cell line was generated based on our previous work done in other cell lines^11^ to serve as a benchmark. Gal8-YFP tracking shows the same overall trend for PPAA in HEK293-T cells, with a strong dose response to PPAA treatment (Figure S6). In comparing the Gal8-YFP system to G8G8 population 4, we observed that the split luciferase G8G8 system better excludes high, toxic doses of PPAA, as occurs at 625 μg/mL (Figure 3). When loss of cellular metabolic activity occurs, the G8G8 system showed a loss of luminescent signal while the Gal-YFP assay shows a very high Gal8-YFP response. While the Gal-YFP assay must be used with cytotoxicity measurements in parallel, the G8G8 system gives a positive response only on live and metabolically active cells due to the ATP-dependence of firefly luciferase. Another advantage of the G8G8 system over the Gal-YFP in this study was that it was more sensitive in detecting statistically significant endosomal disruption at 20 μg/mL PPAA (p < 10^−4^), whereas Gal8-YFP recruitment tracking failed to reveal an effect until higher polymer doses.

**Figure 3.**
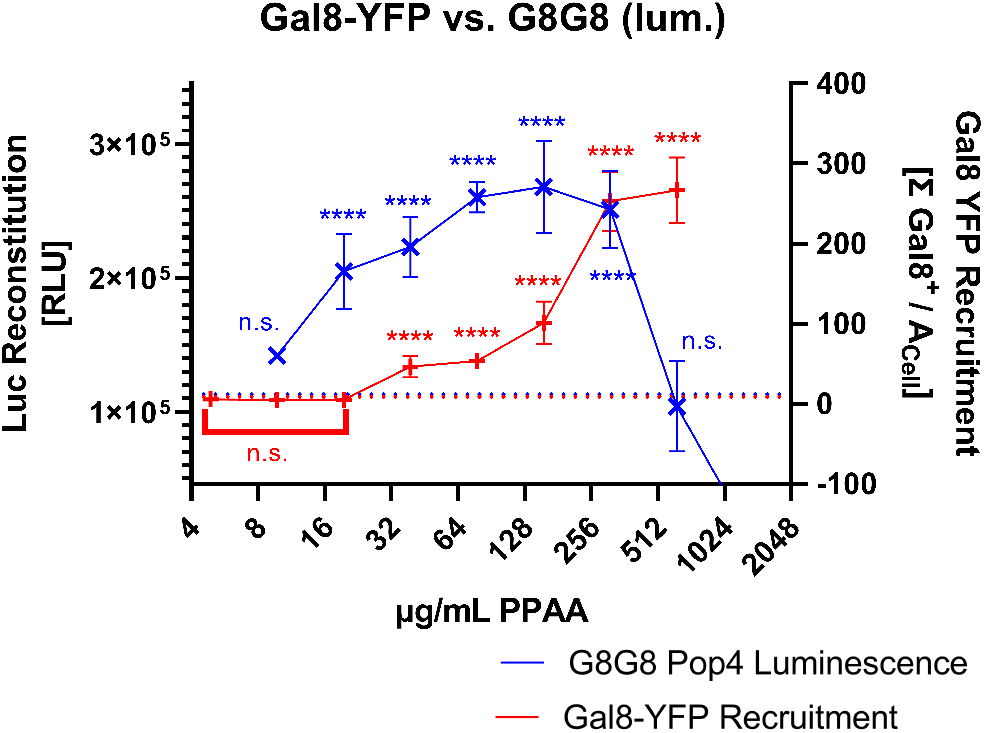
G8G8 cross-validation with the Gal8-YFP assay. Gal8-YFP and G8G8 HEK-293-T cells were treated with a 2-fold dilution series from 1250 to 9.8 μg/mL PPAA for 2 h. G8G8 luminescence data are plotted in blue as × on the left y-axis. Gal8-YFP response data are plotted in red as + on the right y-axis. No treatment control responses were aligned and plotted as dotted line; maximal responses were also aligned to compare responses.

### G8G8 Assay Response to a Broad Set of Transfection Reagents

Next, we assessed the ability of our G8G8 Pop4 cells to detect endosome disruption caused by several other transfection reagents. We treated cells with the commercial cationic transfection lipids DOTAP and Lipofectamine^®^ 2000, as well as the *in vivo* transfection reagent JetPEI^®^. We also screened four additional cationic siRNA delivery 50B polymers that we previously developed^25^, characterized, and screened^11^ for Gal8-YFP recruitment and siRNA delivery efficiency. These polymers have the same composition but varied molecular weights. Molecular weight has a significant correlation to endosome disruption and intracellular bioavailability with this delivery system^11^ (data reproduced as Figure S7 with 50B-S, -L, -2XL, and -4XL having increasing molecular weights). We show that G8G8 cells produce a significant increase in cellular luminescence for all agents tested. DOTAP, JetPEI^®^, and Lipofectamine^®^ 2000 all showed significant increases in luminescence within 2 h of treatment, and, similar to the results with PPAA, allowed for identification of toxicity at the highest doses (Figure 4 A-C). Because DOTAP was formulated in DMSO (100 mg/mL stock in DMSO) and Lipofectamine^®^ 2000 is formulated in ethanol (proprietary concentration), we also ran separate control experiments to show that neither DMSO nor ethanol treatment is sufficient to induce increases in luminescence in this assay format, showing only cytotoxicity at high doses (Figure S8). When we screened four members of our 50B library for G8G8 luminescence, we found that 50B-L, -2XL, and -4XL all showed significant luminescence, while 50B-S did not produce significant luminescence (Figure 4D). This mirrors our earlier bioactivity, TEM, and Gal8-YFP recruitment data for these polymers^11^ (partially reproduced in Figure S7), which showed that the 50B-S polymer did not cause endosomal disruption, did not recruit Gal8, and did not cause siRNA-mediated gene knockdown, in contrast to the larger molecular weight polymers. The assay fully recapitulated the rank-order of endosome disruption previously measured by Gal8-YFP at high doses for the polymer library, where greatest disruption is seen for 50B-2XL and 4XL, followed by - L, with 50B-S showing no activity. DOTAP and JetPEI^®^ showed luminescent responses *in vitro* at manufacturer recommended ranges, which are 50-200 μg/mL and 0.2- 0.4% v/v, respectively. We did not, however, detect endosomal disruption at the supplier-recommended dose of Lipofectamine^®^ 2000 (0.2-0.5%). Further exploration of a time-course following cell treatment with 0.5% v/v Lipofectamine^®^ at 0, 2, 8, and 24 hours revealed that this reagent/dose peaks later, at 8 h for the time points that we measured (Figure S9). We likewise conducted a similar assay with PPAA (Figure S10) at 15, 30, 60, and 120 minutes. This study revealed that high doses initially show high luminescence but cause toxicity at longer timepoints, while lower doses show endosomal disruption that peaks after about 30 minutes but remains strong for at least 120 minutes.

**Figure 4.**
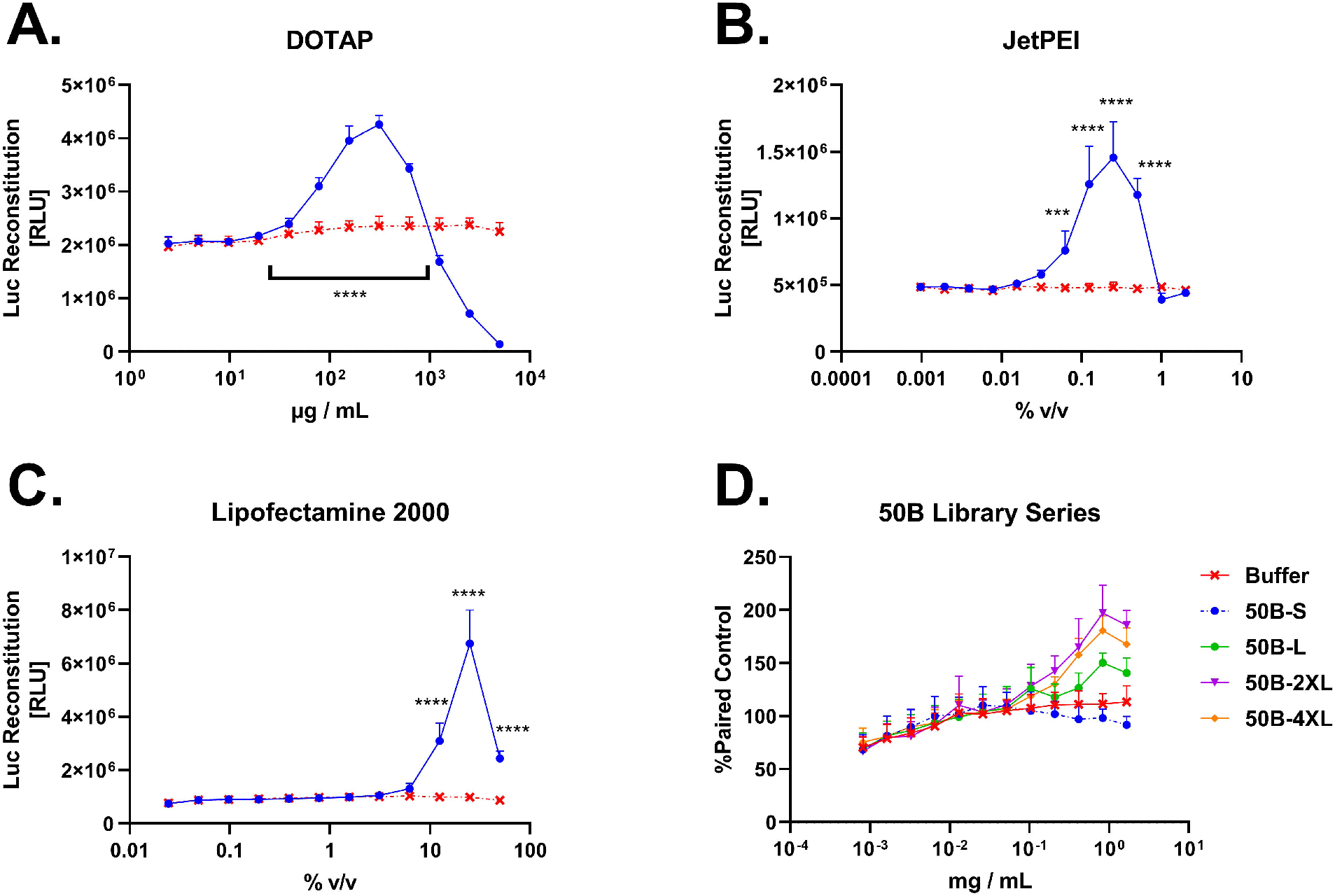
G8G8 Pop4 demonstrates detection of endosomal disruption from several commercial and investigational compounds. (A) HEK-293-T G8G8 cells were treated for 2 h with a 2-fold dilution series of DOTAP lipid in DMSO, starting at 5 mg/mL to 2 μg/mL. Toxicity is observed at high doses until 625 μg/mL, which shows significant luminescent response. Signal returns to baseline at 39 μg/mL DOTAP and below. DMSO alone does not cause luciferase reconstitution (Figure S8). (B) HEK-293-T G8G8 cells were treated for 2 h with a 2-fold dilution series of JetPEI^®^ as formulated by the manufacturer, starting at 2.0% v/v to 0.001% v/v. Baseline response is observed at 2.0% and 1.0%; significant luciferase reconstitution is observed from 0.5% to 0.06% which shows significant luminescent response. Signal returns to baseline at 0.03% and below. (C) HEK-293-T G8G8 cells were treated for 2 h with a 2-fold dilution series of Lipofectamine^®^ 2000 as formulated by the manufacturer, starting at 50% v/v to 0.02% v/v. At 50%, 25%, and 12.5% v/v, significant luciferase reconstitution is observed, which peaks at 25%. Signal returns to baseline at 6.25% and below. (D) HEK-293-T G8G8 cells were treated for 6 h with a 2-fold dilution series of 50B polymers, starting at 1.665 mg/mL to 800 ng/mL. Responses were normalized to same plate buffer readings and plotted. Significant responses were observed for 50B-4XL, 50B-2XL, and 50B-L. 50B-S did not produce a significant response. Significant toxicity was not observed in this dose and time regime. For full statistical analysis for this figure, see Supplemental Tables 3 and 4.

## Conclusions

The development of intracellular-acting large molecule biologic drugs remains limited by cellular barriers to intracellular delivery, especially the plasma and endosomal membranes. While several assays have been developed to assess intracellular drug delivery, assays to specifically measure endosomal disruption are cumbersome, indirect, and susceptible to various artifacts. Here, two first-in-class, genetically-encoded, split luciferase-based, turn on assays are introduced that enable use of luminescence as a rapid and sensitive measurement of endosomal disruption in live cells. Like previous fluorescent Gal8 microscopy assays, these new assays are carrier-and drug-agnostic and do not require labeling of either drug or carrier with tracers, dyes, or synthetic fluorophores which could alter cellular uptake or trafficking. We show that our lead system, G8G8, has significant, dose-dependent luminescent responses to PPAA, Lipofectamine^®^ 2000, DOTAP, JetPEI^®^, and 50B polymers; importantly, this assay is even more sensitive to detecting response to lower concentrations of PPAA than the fluorescent Gal8-YFP assay. We also demonstrate that this assay can be used to assess the kinetics of endosomal disruption, an advantage over many destructive endpoint assays. This assay also has the advantage of having an inherent capacity to rule out cytotoxic treatments because signal generation is dependent on presence of intracellular ATP. In all, the systems presented here are first-in-class turn on luminescence-based assays that directly, rapidly, and quantitatively measure endosomal disruption in live cells without requiring addition of labels to carriers or cargo.

## Materials and Methods

### Plasmid Design

Lentiviral transfer plasmids were designed *in silico* and generated by VectorBuilder (Chicago, IL, USA). The G8C2 system was designed as follows: the first ORF contained a human eukaryotic translation elongation factor 1 alpha1 short form (EFS) promoter driving an mRNA transcript containing NLuc398 (Luciferase AA 1-398), a “GGGGS” triplet spacer (3xG4S), and full length Gal8 (hLGALS8, NM_201545.2), followed by full length human CALCOCO2 (NM_001261390.1), 3xG4S, CLuc394 (Luciferase AA 394-550), then a stop codon. The second ORF consisted of a cytomegalovirus (CMV) promoter driving blasticidin-S deaminase. The G8C2 plasmid has been deposited with AddGene.com (ID 128387). A plasmid map is shown as Figure S11.

The G8G8 system was designed as follows: the first ORF contained a human eukaryotic translation elongation factor 1 alpha1 (EF1A) promoter driving an mRNA transcript containing NLuc398 in frame with a 3xG4S spacer to the first 157 amino acids of human Galectin 8, corresponding to the N-terminal carbohydrate recognition domain (G8NCRD) with a stop codon, followed by an internal ribosome entry site derived from the encephalomyocarditis virus upstream of CLuc394, a 3xG4S spacer, followed by the same G8NCRD. The second ORF consisted of a cytomegalovirus (CMV) promoter driving enhanced green protein connected to blasticidin-S deaminase *via* a T2A self-cleaving peptide. The G8G8 plasmid has been deposited with AddGene.com (ID 128388). A plasmid map is shown as Figure S12.

The NLuc and CLuc degree of luciferase fragment overlap (4 AA) and split point (1 - 398; 394 - 550) were chosen based on work by Paulmurugan and Gambhir,^19^ which showed this design to produce strong luminescence upon protein-protein interaction.

### Lentiviral Preparation

Psuedotyped lentiviral particles (LV) were generated using transfer plasmids (above), pCMV delta R8.2 (AddGene Cat. No. 12263), and pMD2.G (AddGene Cat. No. 12259). Such that a final volume of 600 μL was reached, the following components were added, in order, to a 15 mL polypropylene conical tube to yield a transfection mixture: Opti-MEM media (Gibco, Cat. No. 31985062), 0.6 μg pMD2.G, 3.0 μg pCMV delta R8.2, 6.0 μg transfer plasmids, and 42 μL FuGENE 6 (Promega, Cat. No. E2691). The tube was gently flicked to mix the plasmids before and after the addition of FuGENE 6. The transfection mixture was added dropwise to a T-75 flask at approximately 50% confluency of HEK-293-T (American Type Culture Collection Cat. No. CRL-3216) cells in 11.0 mL DMEM/F-12 supplemented to 10% FBS without antibiotics. After 18 h incubation, the media on the HEK-293-T cells was exchanged for DMEM supplemented with 10% FBS and 1% penicillin-streptomycin. At 24 h, 48 h, and 72 h after this media change, the virus-containing supernatant was harvested, syringe filtered (0.45 μm, nylon), and concentrated from ~11 mL to ~250 μL with Amicon Ultra-15 centrifugal filtration unit (100 kDa nominal molecular weight cut off). The LV were then either used immediately or aliquoted and frozen for later use.

### Cell Line Generation

A 2-fold dilution series of LV was made in serum free, HEPES supplemented DMEM, starting with undiluted viral concentrate (2^0^) and ending with 2^−11^ viral supernatant. To 12 wells of a 96-well plate, 100 μL of 10^5^ cells / mL were added. Then, 100 μL from the dilution series was added to the plate of cells. After 48 h in which the cells adhered and proliferated, the media was changed and selection began, using DMEM supplemented with 10% FBS and 5 μg/mL blasticidin. Cells were monitored by phase contrast microscopy; as wells reached confluence, cells were washed twice with PBS −/−, trypsinized with a minimal volume of 0.25% trypsin, diluted with serum containing media, and transferred without centrifugation to progressively larger vessels. Media was changed every 48 – 72 h. As each T-75 flask reached ~80% confluence, half the cells were aliquoted and stored cryogenically in a 90:10 v/v mixture of FBS and DMSO and further expanded using T-175 plates. Cells were periodically monitored throughout this process for EGFP expression *via* microscopy or for luciferase expression by IVIS imaging. HEK 293-T cells stably expressing G8G8 or G8C2 are available upon request from the corresponding author. Cells originating from the first five dilutions were denoted Population 1 through Population 5 and used herein.

### Luciferase Measurements

Luciferase measurements were taken on live cells by exchanging media with media supplemented with 10% FBS, 150 μg/mL D-luciferin, and 25 mM HEPES. Cells were imaged after 3-5 minutes of incubation at 37 °C in clear bottom, black wall, 96-well plates in an IVIS Lumina Imaging system (Xenogen Corporation, Alameda, CA, USA) with 60 s exposure. Luminescence was monitored for up to 20 min. ROIs were drawn and average photon flux was calculated and exported.

### Gal8-YFP Measurements

Gal8-YFP measurements were conducted as previously reported^8^ with two differences. First, HEK293-T were used instead of MDA-MB-231 to generate stable Gal8-YFP expressing cells. Second, recruited Gal8-YFP was normalized to total cellular area rather than cell number by quantifying cellular area by thresholding pixels above background. HEK-293-T Gal8-YFP cells and Gal8-YFP retrovirus concentrates are available upon request from the corresponding author. A PiggyBac compatible Gal8-GFP has also been made available by the Jordan Green lab at AddGene (ID 127191).^10^

### Cellular Viability Measurements

HEK-293-T G8G8 cells were seeded into a 96 well plate and treated with a 2-fold dilution series of PPAA, starting with 625 ug/mL. After 2 hours, the PPAA media was removed and cells were washed once with DMEM supplemented with 10% FBS. Then, media was changed to DMEM supplemented to 10% FBS and 25 ug/mL resazurin salt. Plate absorbance was read on a TECAN Infinite M1000 Pro plate reader in a continuous kinetic cycle.

### Lipofectamine Kinetic Studies

HEK-293-T G8G8 cells were seeded in a 96 well plate and treated with 0.5% v/v Lipofectamine 2000 in serum free DMEM at 2, 8, or 24 hours before reading. After the specified amount of time, the media was replaced with luciferin-containing media and luminescence was measured on the IVIS.

### PPAA Kinetic Studies

HEK-293-T G8G8 cells were seeded in a 96 well plate and treated with a 2-fold dilution series of PPAA in serum-free DMEM starting with 625 ug/mL. The PPAA treatment was left on the cells for 15, 30, 60, or 120 minutes, after which the media was replaced with luciferin-containing media and luminescence was measured on the IVIS.

### Data Processing and Statistical Methods

Plots, regressions, and statistics were generated using Prism GraphPad 8. For the G8C2, G8G8, and Gal8-YFP systems, PPAA versus vehicle dose response curves were analyzed using ordinary two-way ANOVA with the following settings: no sample matching, column factor of putative endosome disrupting agent versus buffer, row factor of dose, with post-hoc testing to compare dose matched response of agent versus buffer. Sidak’s multiple comparison correction was used to calculate a multiplicity adjusted P value. The family-wise significance and confidence rate was set to 0.05, or a 95% confidence interval. For figure S5, ordinary linear regression was performed. For figure S9, ordinary one-way ANOVA was performed with multiple comparisons testing against control. Throughout the manuscript, asterisks are used *per* the GraphPad Prism style: * for p < 0.05, ** for p < 0.01, *** for p < 0.001, and **** for p < 0.0001.

#### Material Availability

Plasmids encoding the G8C2 (plasmid ID 128387) and G8G8 (plasmid ID 128386) systems are available from AddGene.com to nonprofit users. For requests of cell lines, please contact the corresponding author.

## Supporting information

Supporting Tables (Supplementary Statistics)

## Supporting Information

Supporting information in the form of several additional figures and supplementary statistics are available for free in the supporting information online at the publisher’s website.

## Acknowledgements

This work was supported by the United States National Institutes of Health grants R01HL122347, R01EB019409, and R01CA224241 to CLD.

---

**Figure S1.**
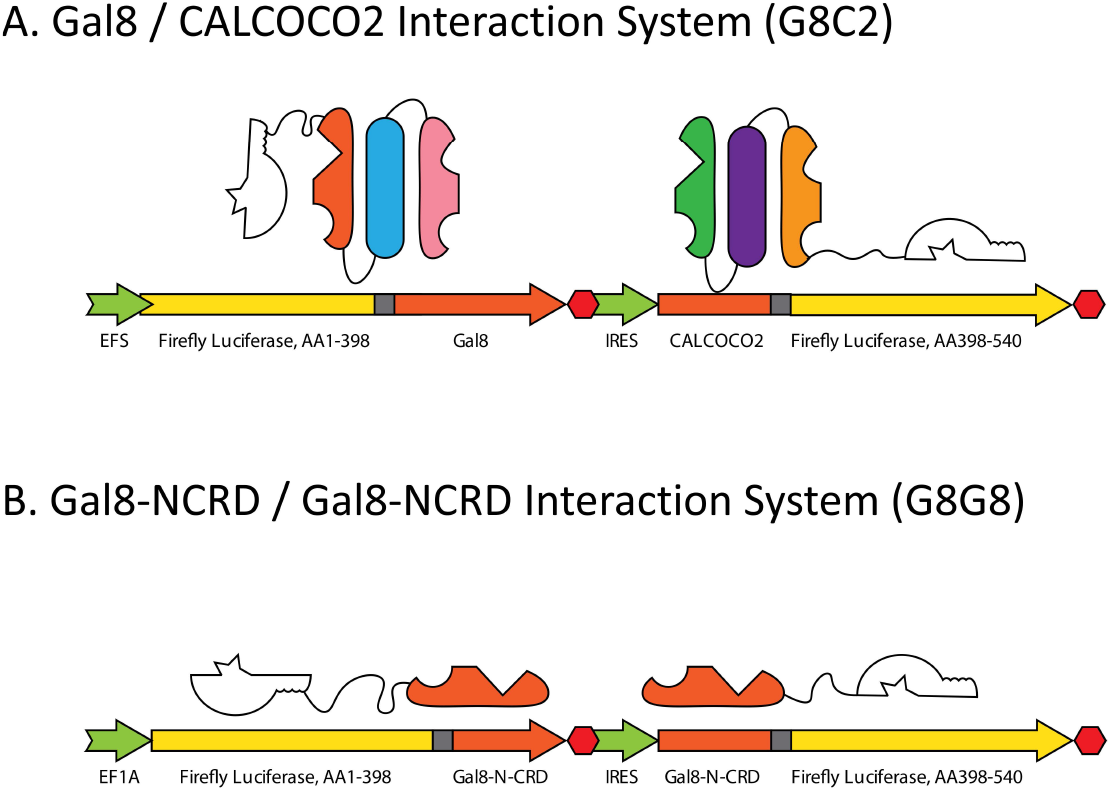
Graphical overview of the G8C2 and G8G8 transcripts. A. From a single mRNA transcript driven by the EFS promoter, two proteins are expressed. The first is NLuc-GGGGSx3-Gal8 and the second is CALCOCO2-GGGGSx3-CLuc, connected by an internal ribosomal entry sequence. B. From a single mRNA transcript driven by the EF1A promoter, two proteins are expressed. The first is NLuc-GGGGSx3-G8NCRD and the second is G8NCRD-GGGGSx3-CLuc, connected by an internal ribosomal entry sequence.

**Figure S2.**
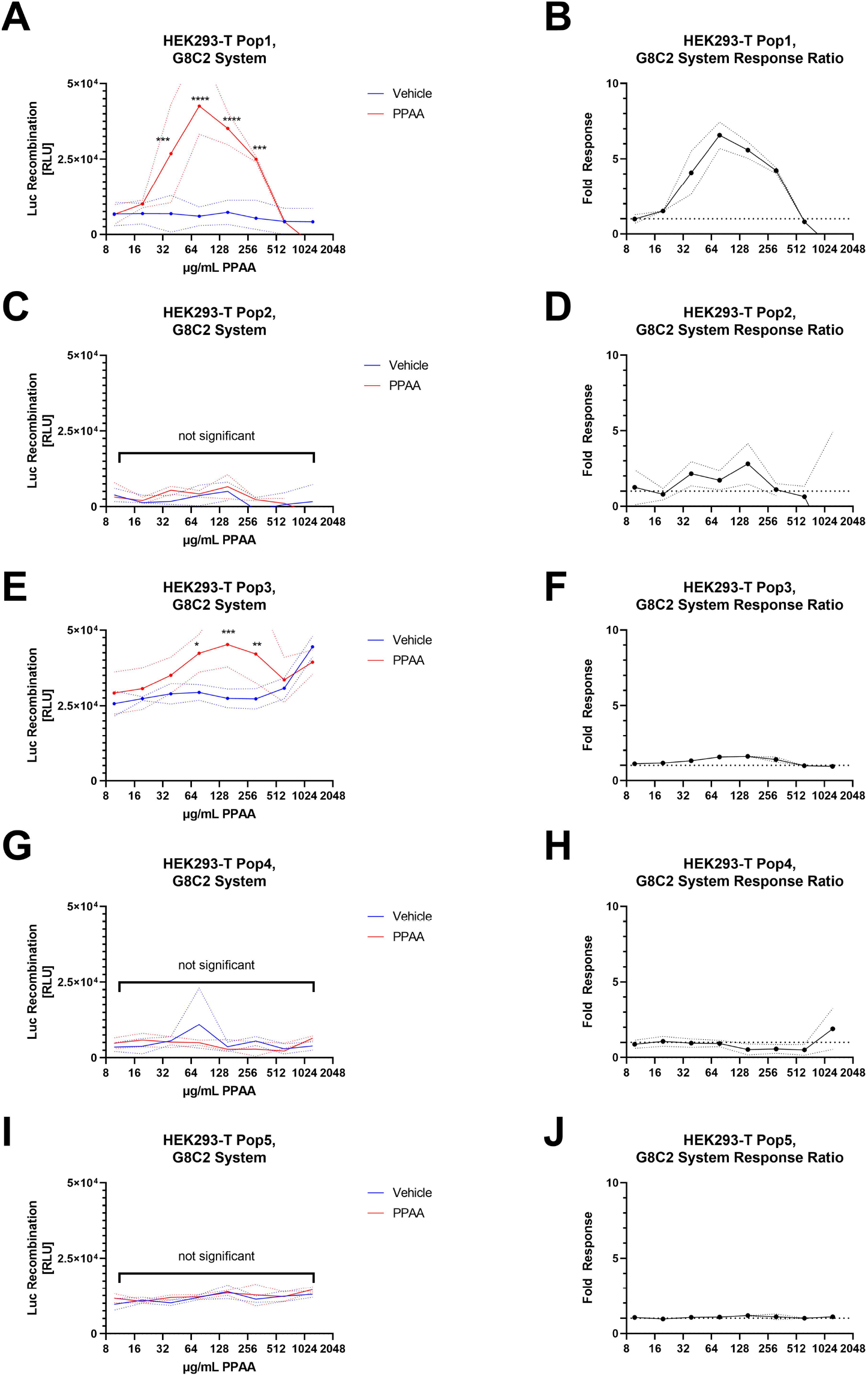
G8C2 cell population screen with PPAA. A, C, E, G, I: G8C2 cells were treated with a 2-fold serial dilution of PPAA for 2 h, media exchanged, and luminescence measured. p values are indicated as: p < 0.05, *; < 0.01 **; < 0.001 ***; < 0.0001 **** B, D, F, H, J: luminescent response was normalized to vehicle values and plotted as fold response versus PPAA concentration

**Figure S3.**
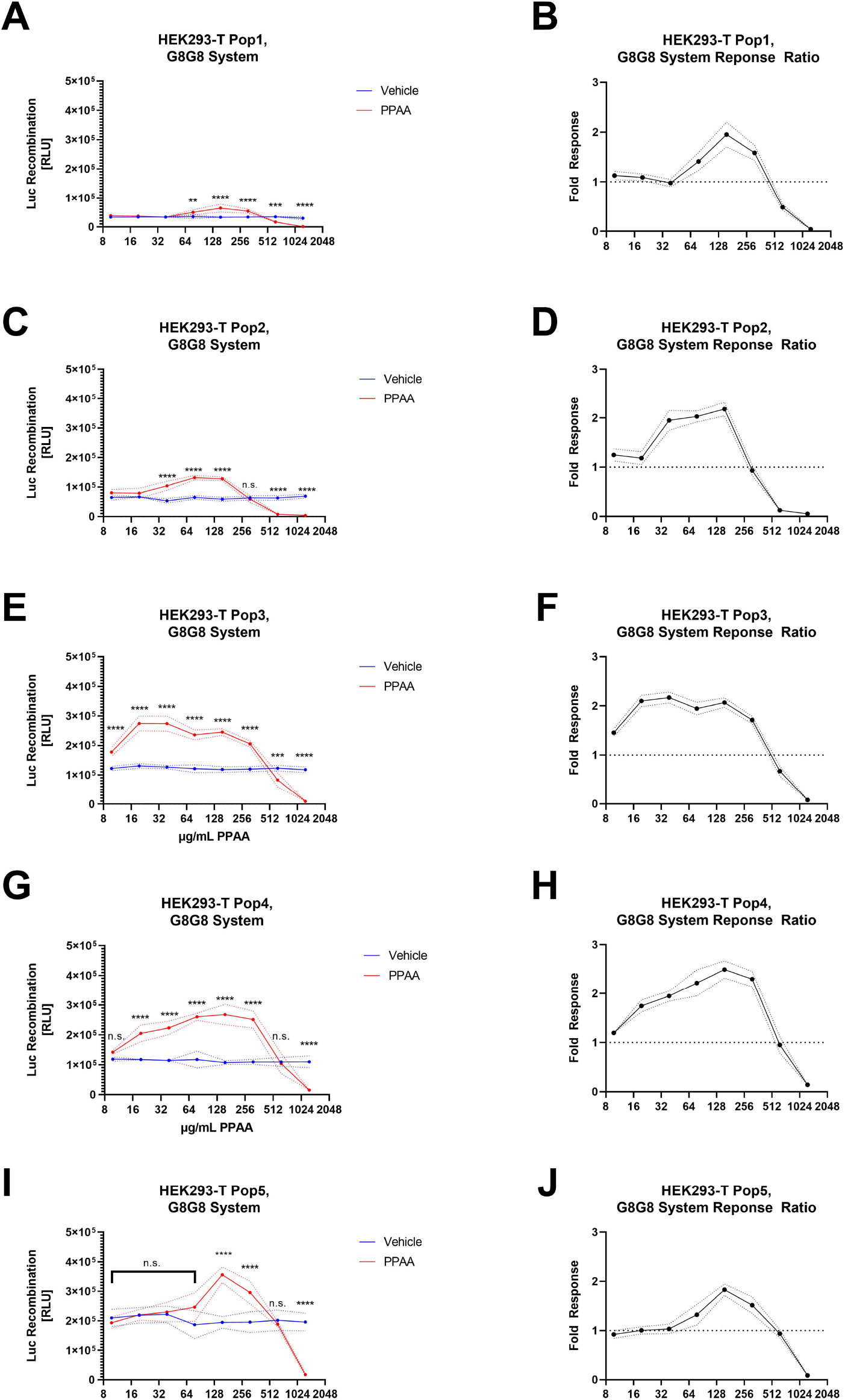
G8G8 population screen with PPAA. A, C, E, G, I: G8G8 cells were treated with a 2-fold serial dilution of PPAA for 2 h, media exchanged, and luminescence measured. p values are indicated as: p < 0.05, *; < 0.01 **; < 0.001 ***; < 0.0001 **** B, D, F, H, J: luminescent response was normalized to vehicle values and plotted as fold response versus PPAA concentration

**Figure S4.**
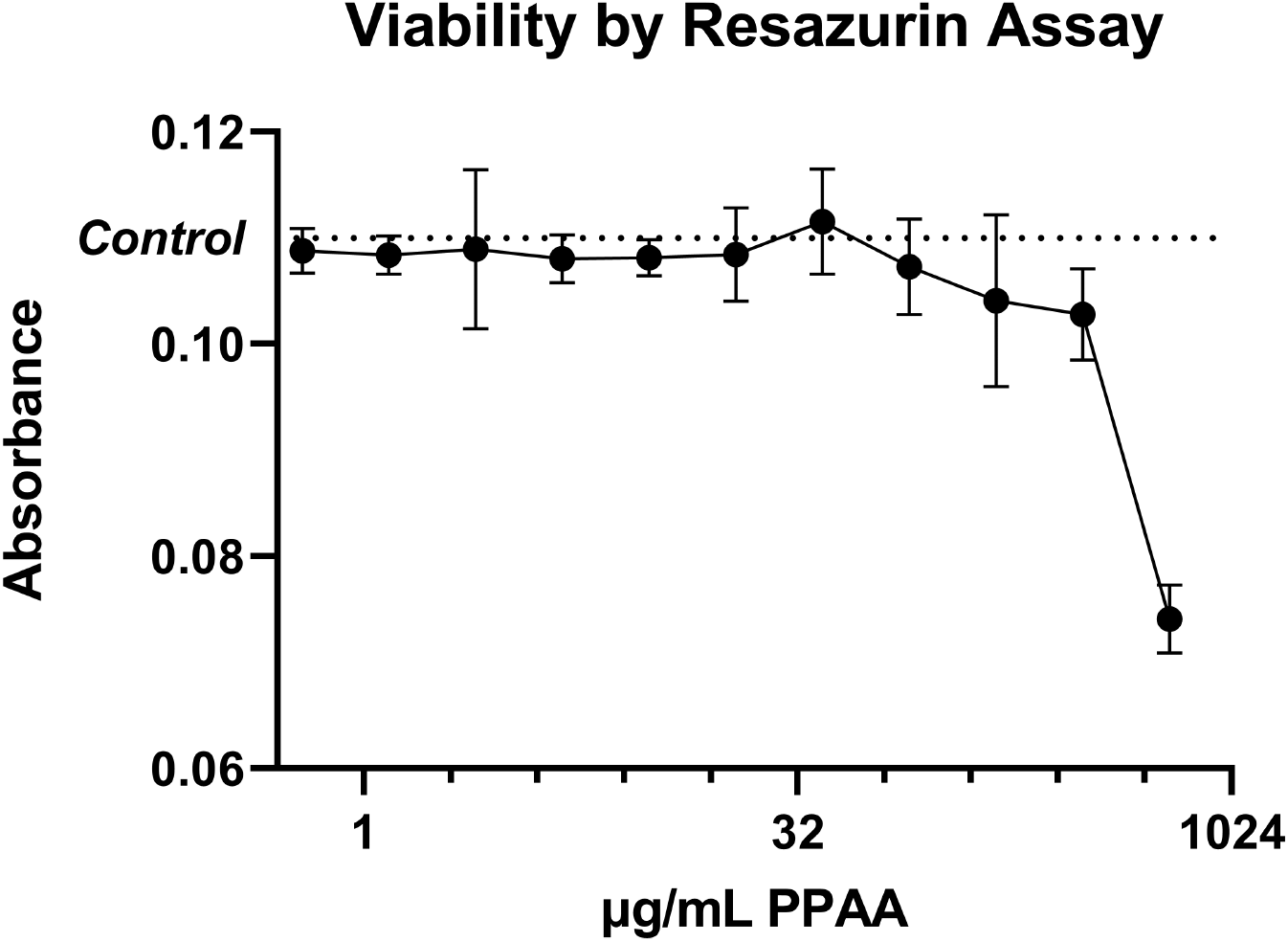
HEK293-T treated PPAA viability screen. G8G8 HEK-293-T cells were treated with a 2-fold dilution series from 1250 to 9.8 μg/mL PPAA for 2 h. Viability was assessed with the resazurin reduction assay. A notable loss of viability is observable at high doses. The average no treatment control signal is plotted as a dotted line.

**Figure S5.**
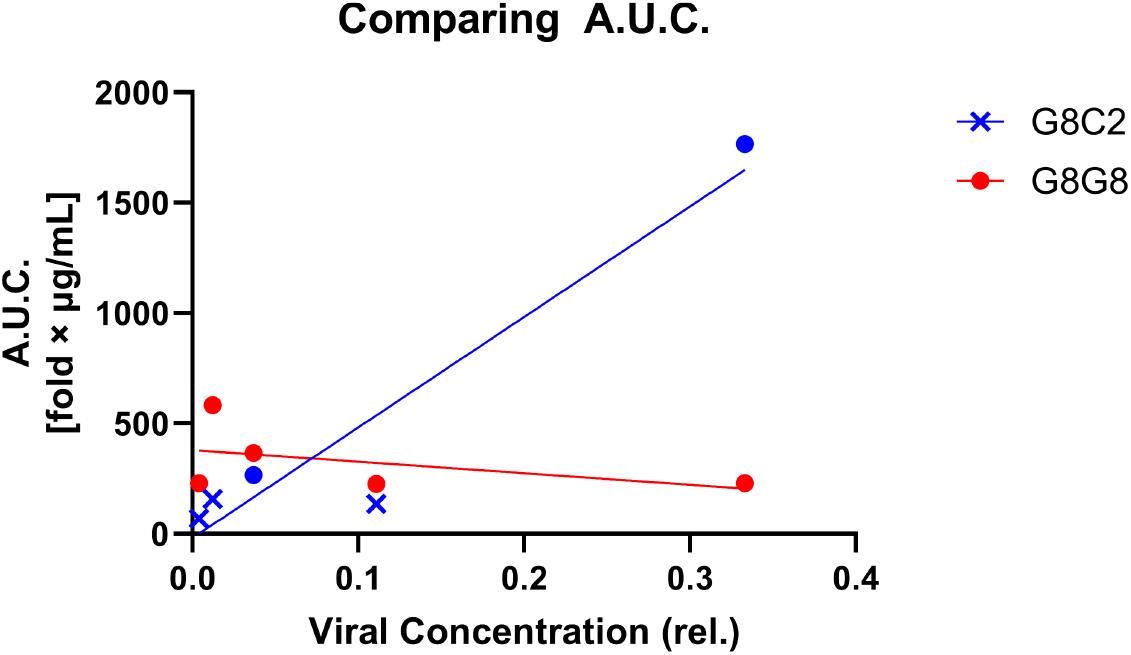
AUC analysis of G8C2 and G8G8 populations. The area under the curve for each PPAA endosome disruption experiment was calculated and plotted against the viral dilution factor to check whether MOI had any effect on system responsivity. The G8C2 system had a statistically significant correlation between MOI and AUC (R^2^ = 0.90, “Significant” non-zero slope), while G8G8 did not (R^2^ = 0.21, “not significant”). Datapoints correspond to clones 1-5 for each system, counting from the right, i.e., the rightmost points are Pop1 and the leftmost points are Pop5. Datapoints are plotted as closed circles if at least one dose of PPAA produced a significantly different response from control and as an x if no dose of PPAA produced such a response.

**Figure S6.**
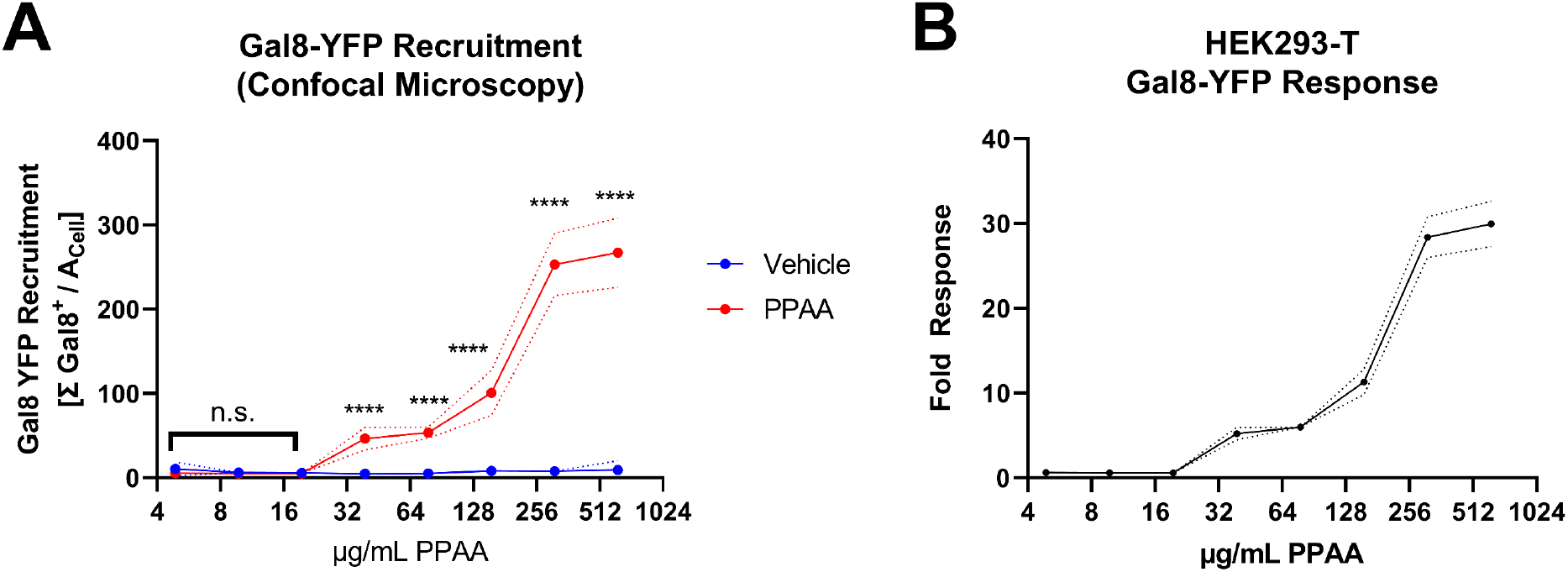
HEK293-T cell line establishment Gal8-YFP dataset. A. Gal8-YFP cells were treated with a 2-fold dilution series from 1250 to 9.8 μg/mL PPAA for 2 h. Gal8-YFP response data are plotted in red and buffer vehicle treated cells are plotted in blue. Significant differences were observed at the five highest doses. B. The Gal8 response was normalized to the average buffer response and plotted as fold response

**Figure S7.**
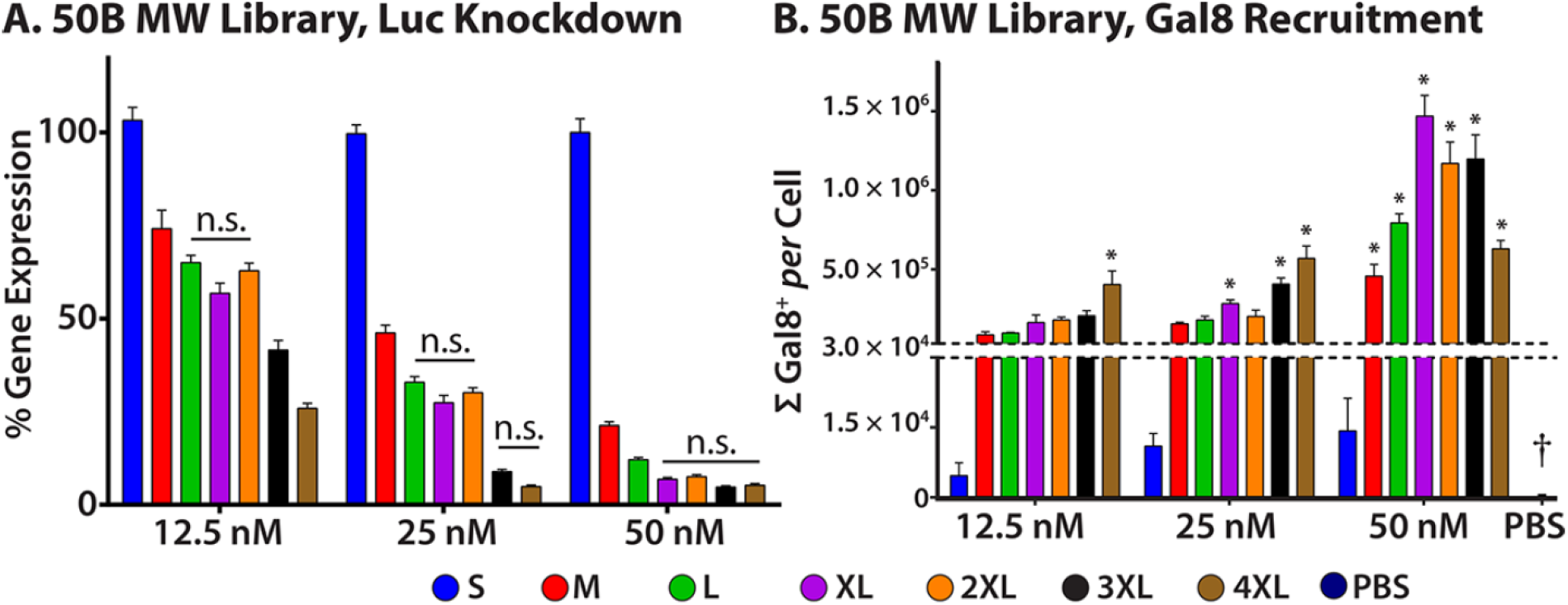
Previously reported 50B polymer bioactivity (gene knockdown) and Gal8-YFP recruitment data for polymers with varied molecular weight. A. 50B shows dose-dependent anti-luciferase siRNA mediated gene knockdown for 50B-M, L, XL, 2XL, 3XL, and 4XL, but not 50B-S polymers. Enhanced siRNA delivery is mediated by enhanced endosome disruption and results in reduced luciferase expression. B. MDA-MB-231 Gal8-YFP cells were treated with noted polymers. Study revealed that Gal8 recruitment increases with increasing siRNA dose and increasing polymer MW. PBS-treated cells had near-zero response, highlighted with a dagger. Single asterisks indicate polymers identified as significant hits by Dunnet’s comparison test. These data demonstrate that 50B-S does not induce endosomal disruption, while 50B-L, 2XL, and 4XL, used in Figure 3D, cause significant endosomal disruption. Reprinted with minor modifications for clarity with permission from our recent paper, Kilchrist et al., ACS Nano 2019, 13, 1136-1152. https://pubs.acs.org/doi/abs/10.1021/acsnano.8b05482 Copyright 2019 American Chemical Society.

**Figure S8.**
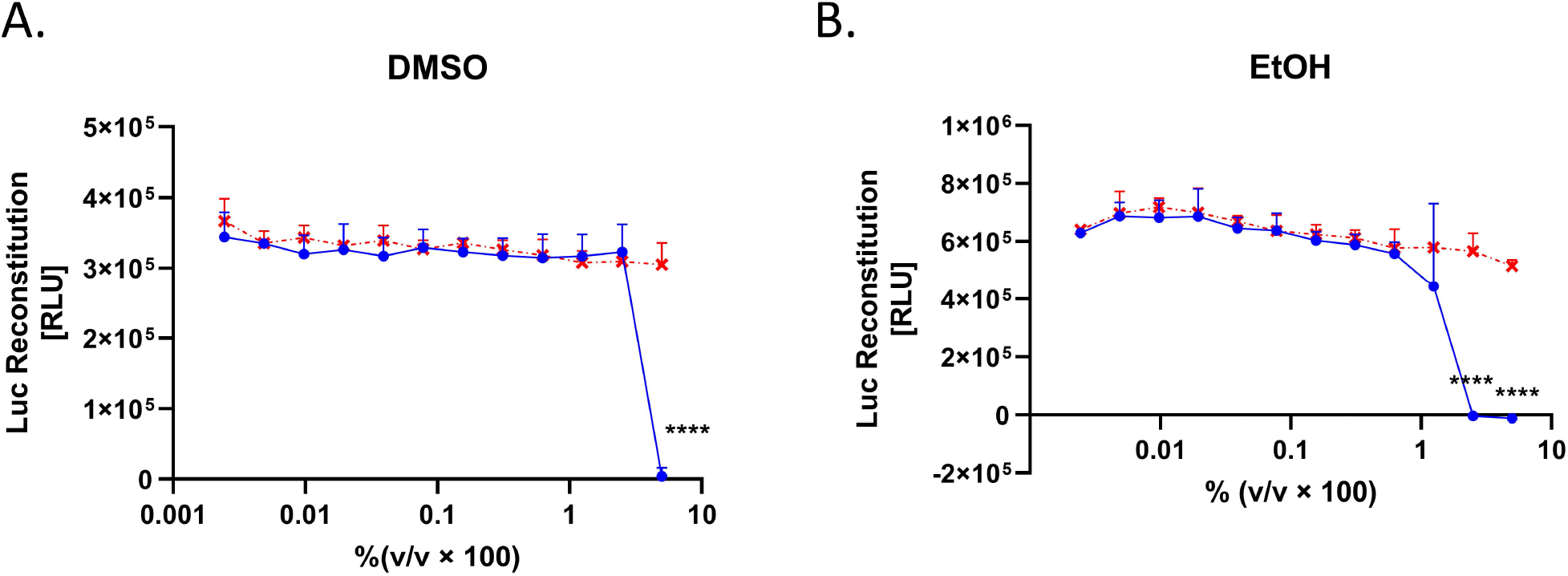
DMSO and ethanol are not sufficient to induce G8G8 response. HEK-293-T G8G8 cells were treated with a 2-fold dilution series of DMSO (A) or ethanol (B) from 5% to 0.002% v/v. No increase in luminescence is detected due to either of these solvent components, which were used in formulating DOTAP (at 100 mg/mL) or Lipofectamine^®^ 2000 (at proprietary concentration). Toxicity is observed for 5% DMSO and 5% and 2.5% ethanol.

**Figure S9.**
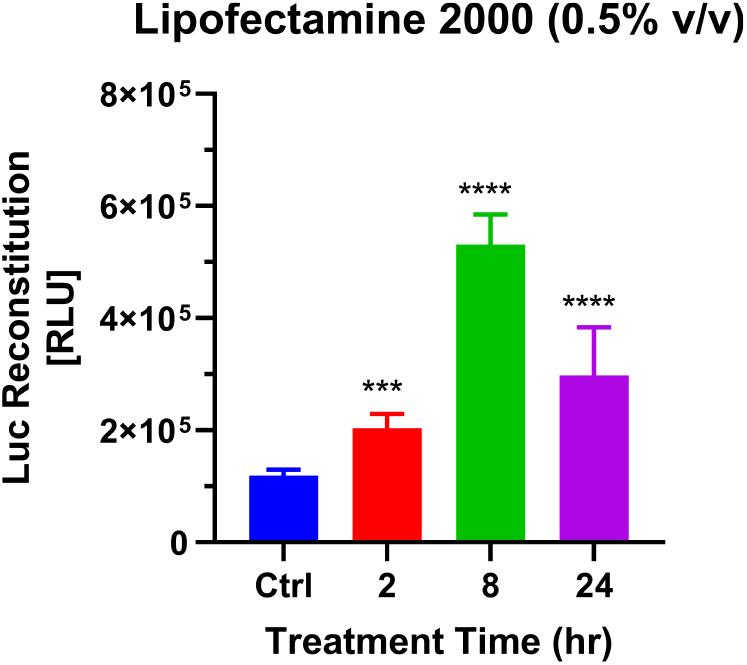
Lipofectamine^®^ 2000 time course of G8G8 split luciferase signal. To resolve the apparent mismatch between endosome disruptive Lipofectamine^®^ 2000 concentrations observed in Figure 3A (50, 25, and 12.5%, but not ≤6.25%), we used the manufacturers recommended concentration, 0.5% v/v in a time course study to resolve this discrepancy. G8G8 HEK-293-T cells were treated with 0.5% Lipofectamine^®^ 2000 for 2, 8, or 24 hr and analyzed for luciferase reconstitution. Significant increases in luminescence relative to control were observed for all timepoints, with maximal response at 8 hr post treatment.

**Figure S10.**
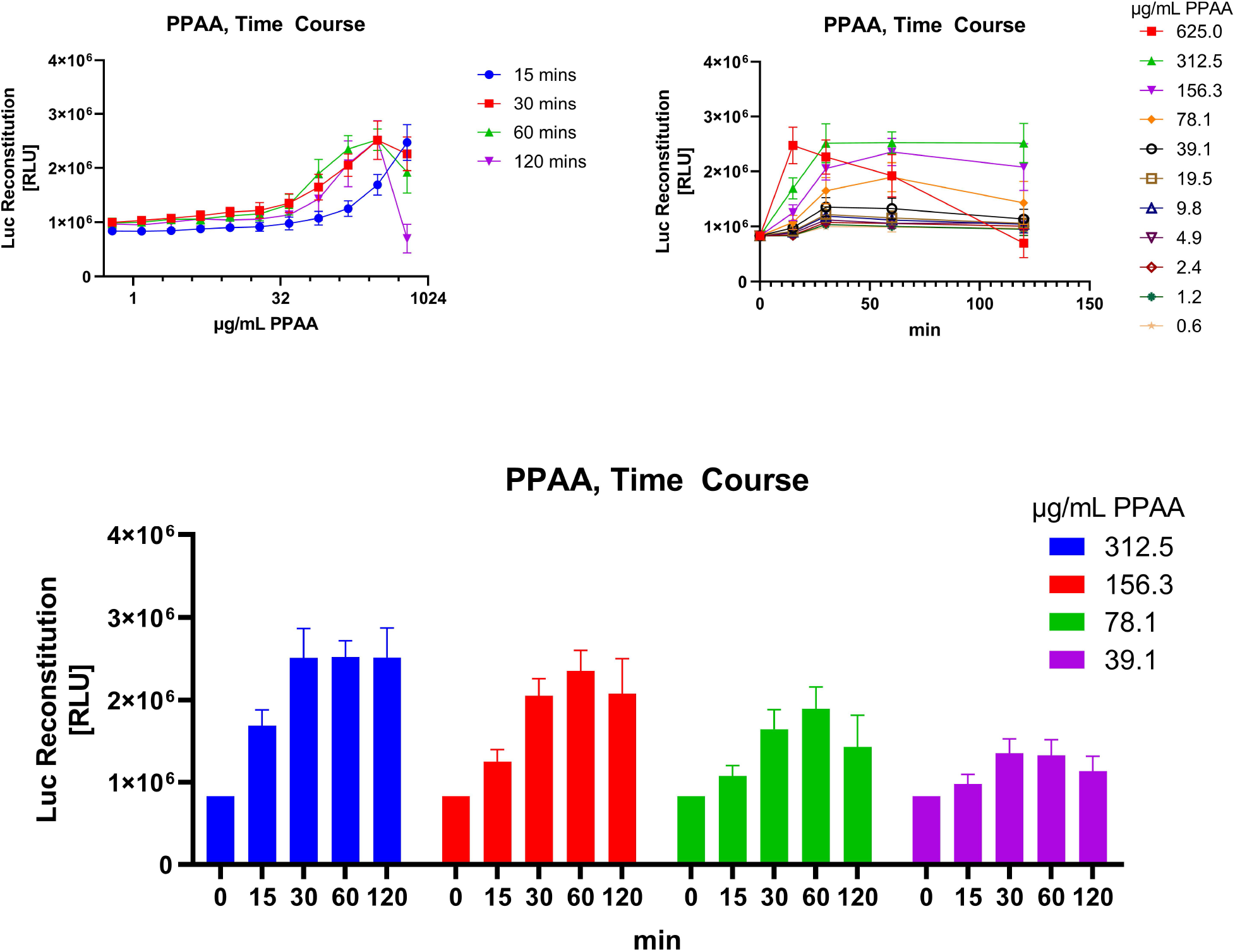
Kinetic testing of PPAA with G8G8 assay system. HEK-293-T G8G8 cells were treated with a 2-fold dilution series of PPAA, starting at 1.25 mg/mL with treatments staggered to provide different contact times. The data from this experiment are plotted three ways. On the top left graph, the dose series for each time point is plotted. On the top right graph, individual doses are plotted as time course series. On the bottom graph, kinetic data for four nontoxic doses are plotted as bar charts. Toxicity is observed at 625 μg/mL at 2 h. These data reveal that endosomal disruption occurs rapidly within 15 minutes for the highest dose, but this dose causes toxicity by 2 hours. More moderate doses show robust endosome disruption without substantial toxicity within the timeframe observed.

**Figure S11.**
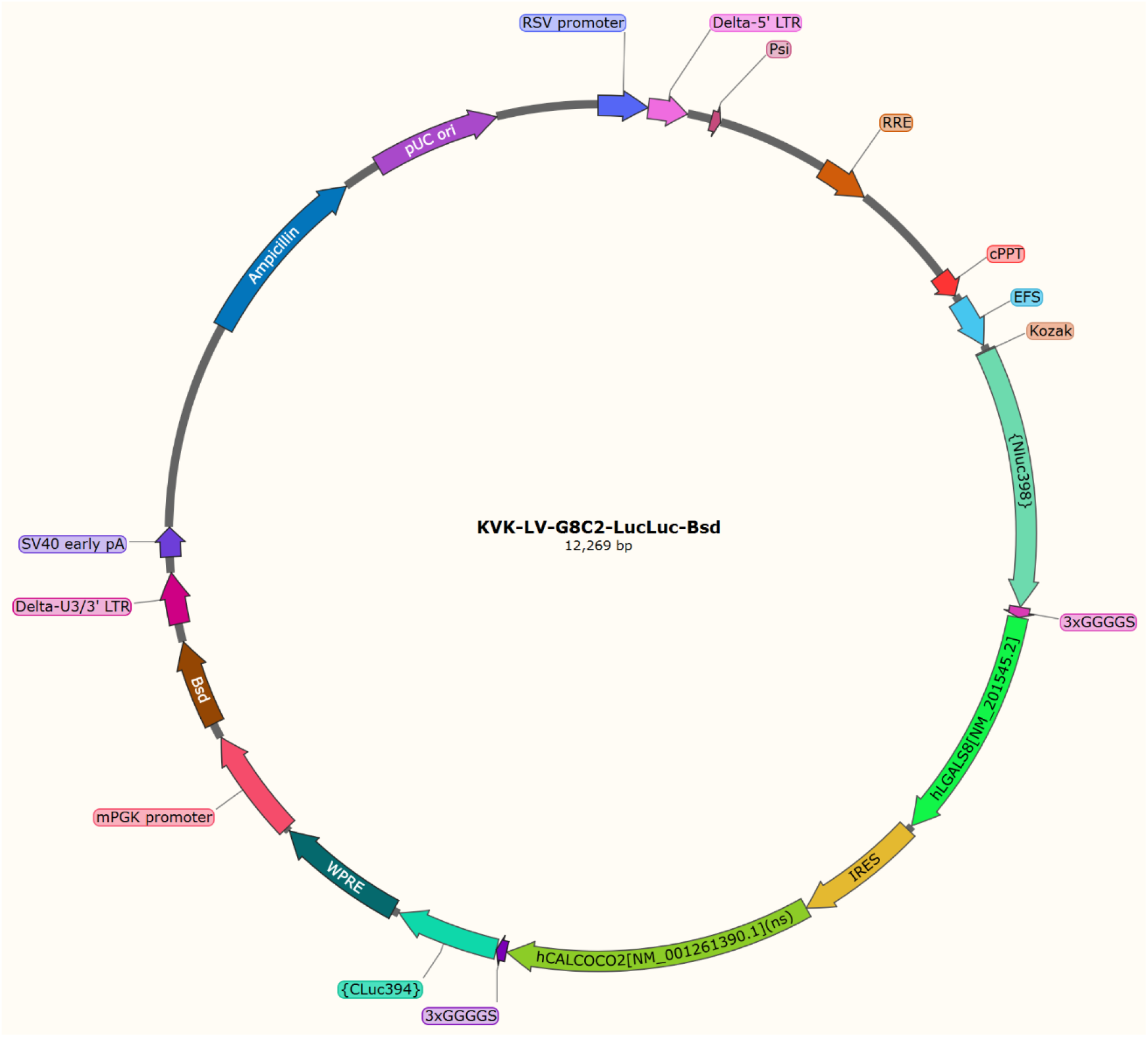
G8C2 Plasmid Map.

**Figure S12.**
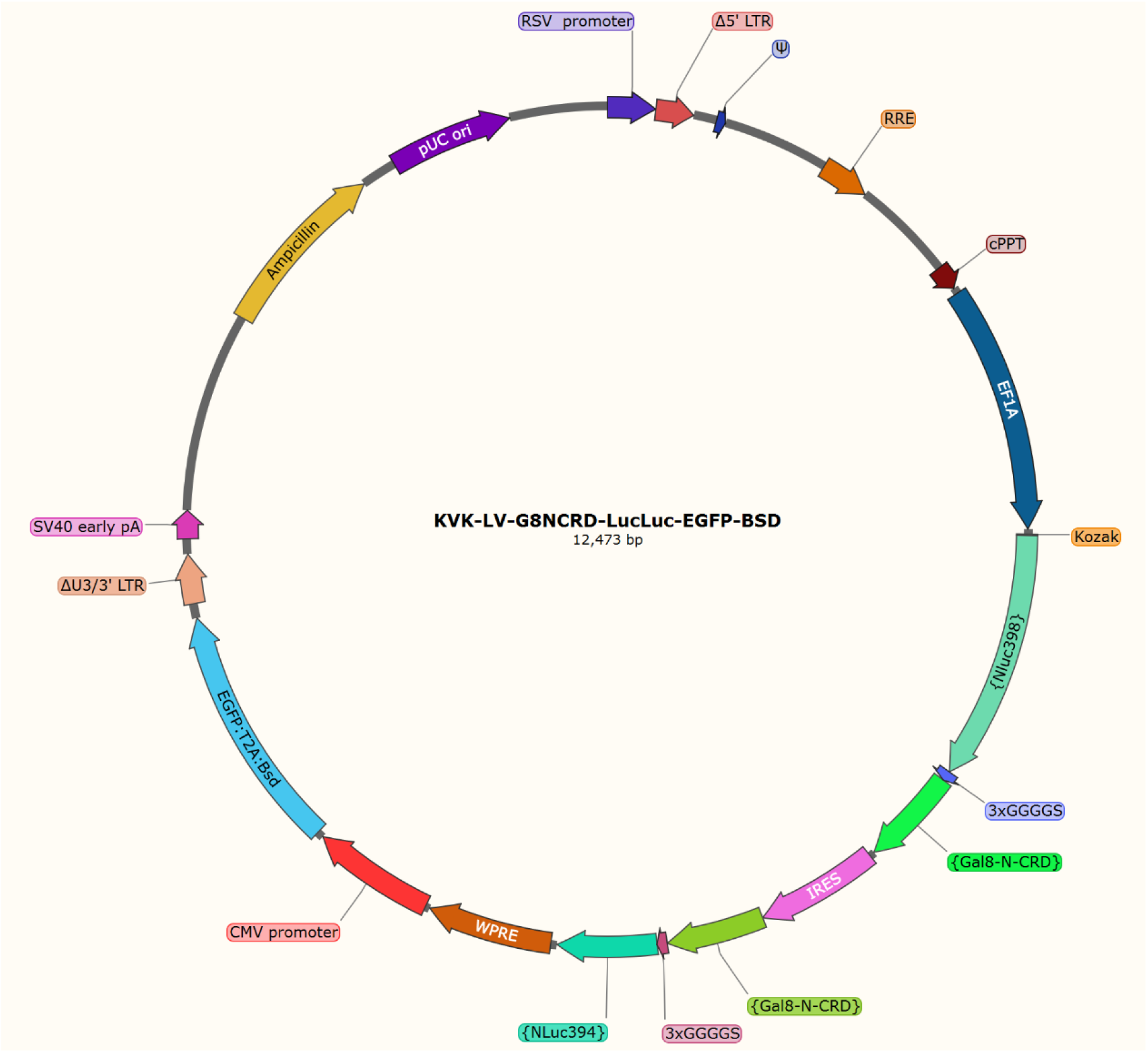
G8G8 Plasmid Map.

